# Wdr5, Brca1 and Bard1 link the DNA damage response to the mesenchymal-to-epithelial transition during early reprogramming

**DOI:** 10.1101/421016

**Authors:** Georgina Peñalosa-Ruiz, Vicky Bousgouni, Jan Patrick Gerlach, Susan Waarlo, Joris V. van de Ven, Tim E. Veenstra, José C.R. Silva, Simon J. van Heeringen, Chris Bakal, Klaas W. Mulder, Gert Jan C. Veenstra

## Abstract

Differentiated cells are epigenetically stable, but can be reprogrammed to pluripotency by expression of the OSKM transcription factors. Despite significant effort, relatively little is known about the cellular requirements for reprogramming and how they affect the properties of induced pluripotent stem cells (iPSC). We have performed high-content screening with siRNAs targeting 300 chromatin-associated factors. We used colony features, such as size and shape, as well as strength and homogeneity of marker gene expression to define five colony phenotypes in early reprogramming. We identified transcriptional signatures associated with these phenotypes in a secondary RNA sequencing screen. One of these phenotypes involves large colonies and an early block of reprogramming. Double knockdown epistasis experiments of the genes involved, revealed that Brca1, Bard1 and Wdr5 functionally interact and are required for both the DNA damage response and the mesenchymal-to-epithelial transition (MET), linking these processes. Moreover, the data provide a resource on the role of chromatin-associated factors in reprogramming and underline colony morphology as an important high dimensional readout for reprogramming quality.

## INTRODUCTION

Somatic cells can be reprogrammed to pluripotency by artificial expression of four transcription factors: Oct4, Sox2, Klf4 and c-Myc (OSKM) (Takahashi and Yamanaka, 2006). With varying efficiency, iPS cells can be derived from a wide variety of cell types and they can differentiate into all cell lineages. Thus, they represent a promising resource for tissue regeneration and disease modeling.

The earliest phase of reprogramming involves dramatic changes in metabolic and cellular processes (Panopoulos et al., 2012; Polo et al., 2012) accompanied by an increase in cell proliferation. Somatic genes are repressed (Maherali et al., 2007; Mikkelsen et al., 2007) and cells undergo MET, a mesenchymal-to-epithelial transition (Li et al., 2010; Samavarchi-Tehrani et al., 2010), leading to the expression of epithelial genes such as *E-Cadherin* (*Cdh1*) and *Epcam*, while mesenchymal regulators (e.g. Snai1/2, Zeb1/2) are repressed (Samavarchi-Tehrani et al., 2010). Subsequently, pluripotency genes carrying active histone marks at regulatory regions are activated (Maherali et al., 2007; Mikkelsen et al., 2007; Polo et al., 2012). At this point, cells have not fully acquired the pluripotency program. These partially reprogrammed intermediates are sometimes referred to as pre-iPS cells (Silva et al., 2008). Late pluripotency markers and endogenous Nanog, Oct4 and Sox2 are activated through a combination of promoter DNA-demethylation (Gao et al., 2013; Meissner et al., 2008) and depletion of repressive histone mark H3K9me3 (Soufi et al., 2012; Sridharan et al., 2013). The majority of the cells seem refractory to reprogramming or are trapped in a partially reprogrammed state, and only a small percentage of cells will successfully progress through all the stages (Polo et al., 2012).

A DNA damage response is important for reprogramming, as the p53 pathway prevents survival of cells with substantial DNA damage (Marion et al., 2009). In agreement with this, DNA repair and recombination proteins are required for reprogramming (Gonzalez et al., 2013; Hansson et al., 2012). Additionally, senescence evokes a DNA damage response and it has been shown to be a barrier for reprogramming (Utikal et al., 2009).

All these events reveal the importance of remodeling the transcriptional program and the chromatin state during reprogramming. Several chromatin-associated proteins that facilitate or block reprogramming have been identified by RNAi (Cacchiarelli et al., 2015; Qin et al., 2014). The activities of the H3K9 methyl transferases Ehmt1/2, Suv39h1/2 and Setdb1 constitute roadblocks of reprogramming (Soufi et al., 2012; Sridharan et al., 2013). In contrast, H3K9 demethylases such as Kdm3a/b and Kdm4c facilitate reprogramming (Chen et al., 2013). In addition, both the repressive Polycomb PRC2 complex (Onder et al., 2012) and the Trithorax SET-MLL methyl transferase complexes act as facilitators. The H3K4 methylation mediated by the SET-MLL complexes primes pluripotency enhancers for activity (Wang et al., 2016) and the absence of their core component Wdr5 abrogates reprogramming (Ang et al., 2011).

Despite the progress that has been made in characterizing the molecular changes during reprogramming, how these dynamic changes are orchestrated is still ill-understood. We have used high-content screening to assess the role of ∼300 chromatin-associated proteins in pre-iPS colony phenotypes during early reprogramming. High-content analysis allows simultaneous measurement of multiple morphological phenotypes. The combination of siRNA screening with high-content microscopy can reveal new associations among pathways (Fischer et al., 2015; Sero and Bakal, 2017). A similar approach has previously been used to define new gene networks involved in the final phase of iPS cells formation (Golipour et al., 2012). We measured over twenty colony-phenotypes, including number of colonies, expression of pluripotency markers and other morphological and textural features, after individual knockdown of 300 chromatin modifiers. Selected hits from the primary screening were subjected to a transcriptome-based secondary screen. We identify several chromatin-associated proteins that act together in the DNA damage response and the MET during early reprogramming to pluripotency.

## RESULTS

### High throughput analysis of the early phase of reprogramming

Reprogramming is associated with major changes in cell morphology, in part due to the MET (Li et al., 2010). Thus, we asked whether chromatin-mediated changes would affect reprogramming efficiency, colony morphology and expression of pluripotency markers. Moreover, we wondered how chromatin-associated factors might work together, as revealed by their similarities in a high dimensional phenotypic space upon knockdown (Mulder et al., 2012; Wang et al., 2012). To define a set of relevant chromatin-associated factors for an siRNA screen (Fig. 1A), we used available expression data (Chantzoura et al., 2015) to select genes with robust expression MEFs or at least 4-fold upregulated expression in reprogramming cells. The custom siRNA library comprised 300 chromatin-associated factors, and for each target, three different siRNA molecules were pooled for transfections (Table S1).

**Figure 1.**
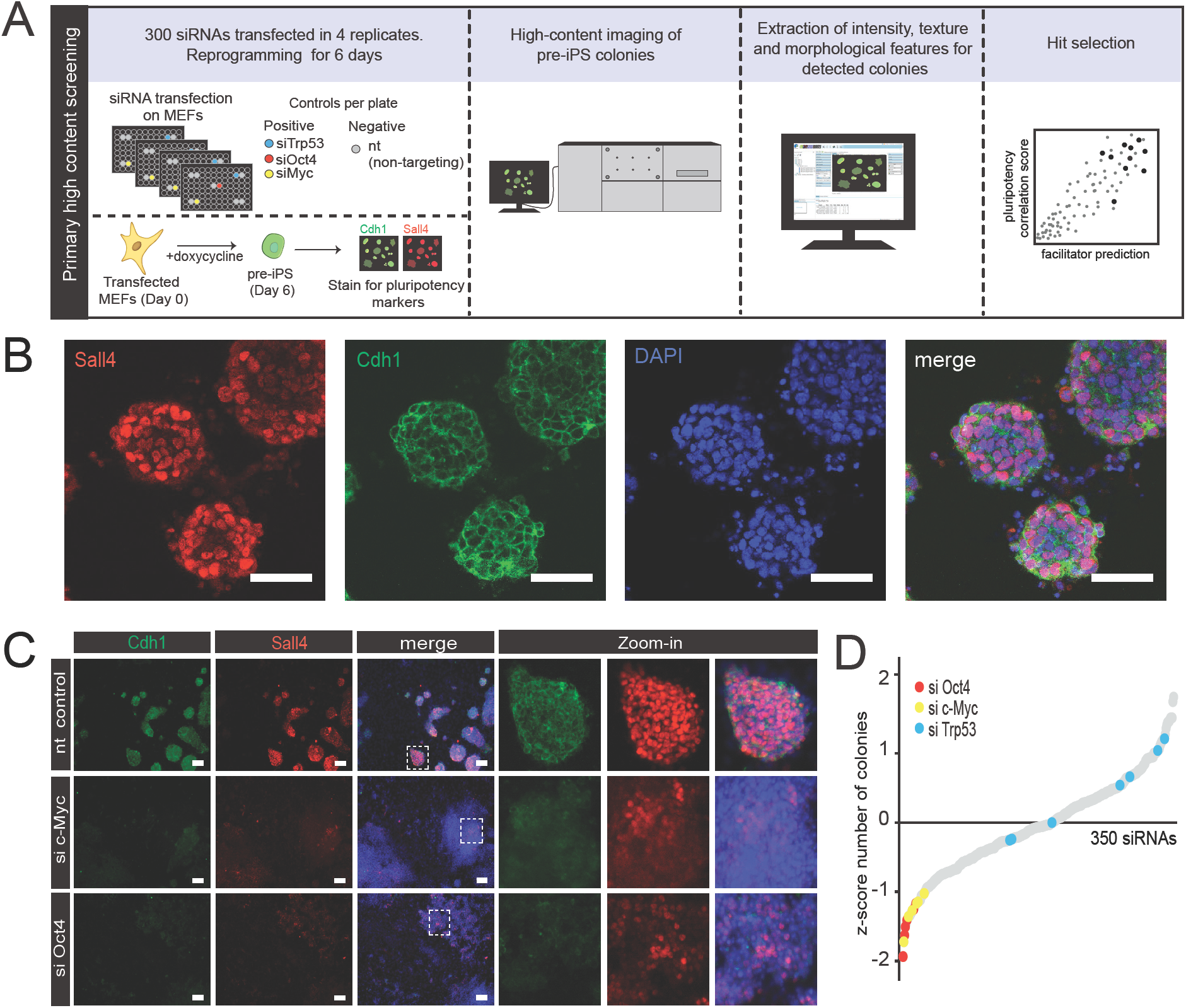
High throughput analysis of the early phase of reprogramming. (**A**) Experimental design of high content imaging siRNA screen. (**B**) Immunofluorescence of pre-iPS colonies at day 6 stained for pluripotency markers Cdh1 and Sall4 with DAPI counterstain. Scale bar represents 50 μm. (**C**) Comparison of colony phenotypes of control, siMyc and siOct4 cells at reprogramming day 6, stained for Sall4 and Cdh1. The scale bar represents 100 μm (left) and the images on the right are a 4x zoom-in from the inset squares on the left. (**D**) siRNAs in the whole screen ranked from low to high z-scores, based on the number of colonies. Positive controls are highlighted in colors. Each siRNA represents the average z-score from four replicates.

We were specifically interested in the early phase of reprogramming, as chromatin is hypothesized to confer epigenetic stability to somatic cells. To test the function of the chromatin-associated genes in early reprogramming, we used a fast and efficient reprogramming system (Vidal et al., 2014), where colonies can be detected after 6 days of reprogramming (Fig. 1B, S1A). These colonies present characteristic round, symmetric morphologies and robust expression of early markers Cdh1, SSEA1 and Sall4, with expression of late markers such as Nanog and Esrrb appearing later (Fig. S1B-D). The specific staining of Cdh1 and Sall4, respectively at the cell surface and in the nucleus, strongly increased between days 3 and 6 (Fig. 1B, S1), representing a suitable readout for the early phase of reprogramming.

The expression of genes was knocked down using siRNAs in mouse embryonic fibroblasts (MEFs) infected with an inducible OSKM-cassette lentivirus. Reprogramming was induced with doxycycline (dox) for 6 days (Fig. S1A). The siRNA library consisted of six 96-well plates, with each plate containing seven non-targeting siRNA (nt) negative controls and three positive controls (siRNA targeting Trp53, Oct4, and c-Myc). The screen was performed in quadruplicate. After six days of reprogramming, samples were fixed, stained for Cdh1 and Sall4 and imaged using an automated high-content microscope. This allowed quantitation of morphology features such as colony size, symmetry and shape, marker intensities, but also texture features, which are a reflection of signal intensity patterns within colonies. After data processing and colony feature extraction, the data were z-score normalized per plate (Bakal et al., 2007) and subjected to further analysis (Fig. 1A).

To test the system, we disrupted reprogramming by knocking down the OSKM factors Oct4 (siOct4) and c-Myc (siMyc). We also knocked down Trp53 (siTrp53), which is expected to enhance reprogramming (Marion et al., 2009). siOct4 and siMyc colonies are flat, irregularly shaped and they show less intense Sall4 and Cdh1 expression compared to the control (Fig. 1C). Likewise, the number of colonies observed in siOct4 and siMyc in our z-score-ranked data was very low (Fig. 1D). The siTrp53 control showed a variable but positive effect on the number of colonies. These data confirm that colony morphology and pluripotency marker expression can be used as readout for a disruption in the early reprogramming network.

### High-content microscopy reveals five major phenotypes of colony formation

The high-content analysis allowed us to measure not only the number of colonies, but also colony features such as the intensity of early pluripotency markers (Sall4 and Cdh1), size, compactness and symmetry, texture and many other morphology features (Table S2). These features constitute a multidimensional phenotypic space for analysis across many conditions or perturbations (Boutros et al., 2015) and the identification of functionally connected genes and processes (Mulder et al., 2012; Wang et al., 2012).

We first defined the set of most discriminating features based on feature-to-feature pairwise correlations (Supplemental Information; Table S3). Using hierarchical (Fig. S2) and K-means clustering (Fig. 2A) we observed five main clusters that display different levels of pluripotency markers, number of colonies, symmetry features (ratio width to length, roundness), STAR morphology features, and textural features (SER, Harlick, Gabor). Cluster 1 knockdowns have few colonies, in addition to low intensities for Sall4 and Cdh1, suggesting a major defect in reprogramming. The majority of nt controls are in cluster 2, which shows a high number of small, round and compact colonies and a robust expression of Cdh1 and Sall4 (Fig. 2A-B). Cluster 3 is quite distinct with fewer, large colonies with low compactness features and detectable Sall4 and Cdh1 expression (Fig. 2A, cf. Brca1 and Wdr5, Fig. 2B). Cluster 4 shows somewhat reduced Sall4 and Cdh1 expression and a reduction in some of the DAPI texture features, but is otherwise relatively normal. Essentially all Oct and Myc controls clustered together in cluster 5, characterized by substantially lower Sall4 and Cdh1 intensities, in addition to irregular, less round and less compact colonies (cf. Ncor1, Fig. 2B).

**Figure 2.**
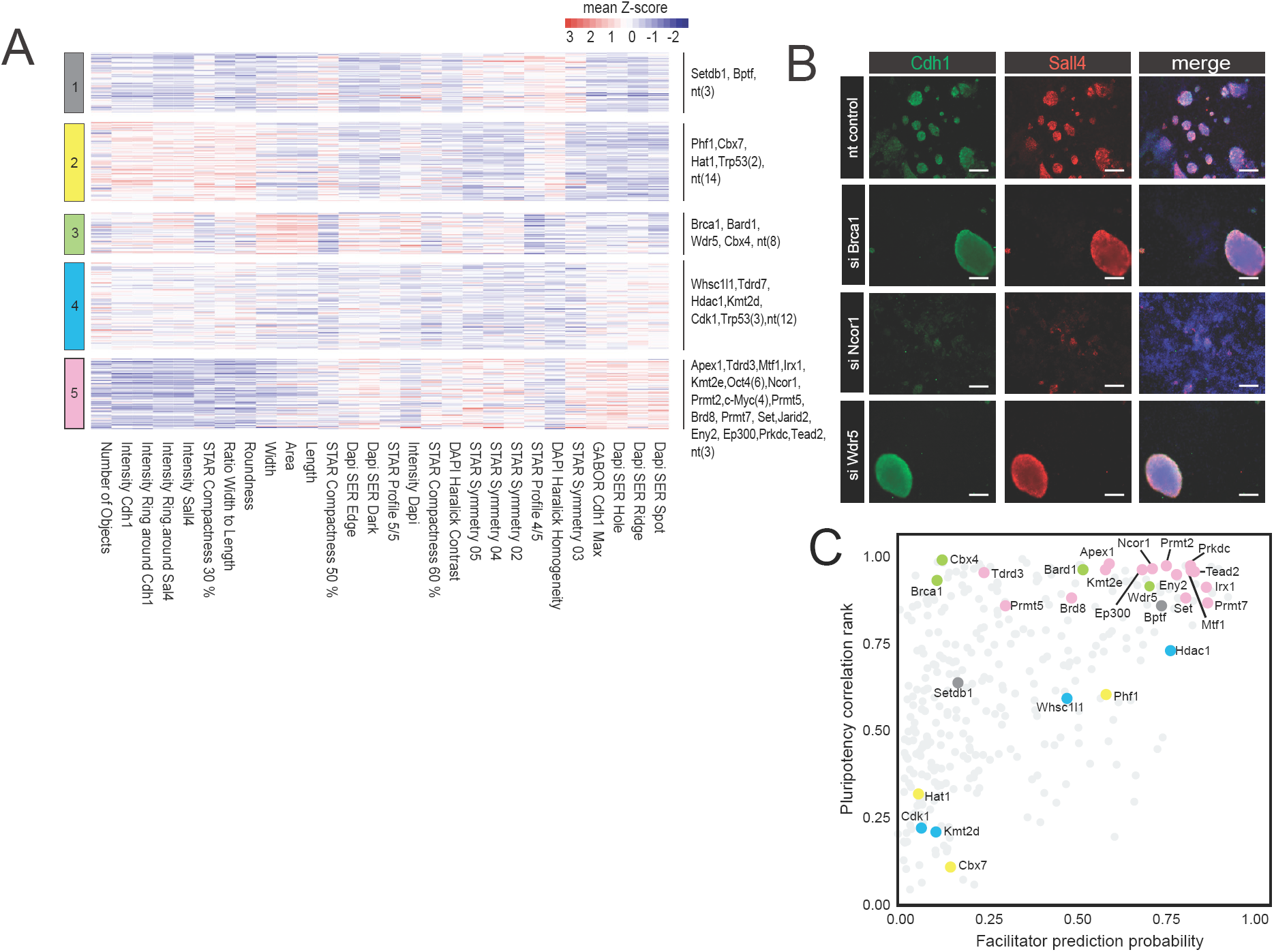
High-content microscopy screen reveals five major phenotypes of colony formation. **(A)** An average Z-score for selected features was calculated from the quadruplicates and represented in a heatmap. Features are clustered by Euclidean distance and rows are clustered by K-means. The bar on the left represents the cluster number and the gene symbols on the right are the hits of the screen and the controls. In brackets are the number of controls in a particular cluster. (**B**) Example images of knockdowns depicting different phenotypes. Scale bar is 200 μm (**C**). Pluripotency-associated hits were selected based on a combination of a probability prediction by machine learning, based on known reprogramming facilitators, and a correlation analysis with the positive and negative controls. Selected top-hits are colored according to the cluster number (panel A, cf. Table S4, Fig. S2).

To provide more insight in the nature of the imaging phenotypes, we compared all knockdowns to each of the positive controls (Trp53, Myc and Oct4) by Pearson correlation, based on high-content features. After ranking all knockdowns according to their combined correlation score (Experimental Procedures), we selected 10 candidates from the top-ranking list (Table S4). Additionally, the high-content data from known reprogramming facilitators present in our library were used to train two independent machine-learning algorithms, in order to predict other potential facilitators (Fig. 2C, Supplementary Information, Fig. S2, Table S4). This approach allowed us to select additional candidates of high, intermediate and low-ranking prediction scores (Fig. 2C). A total of 30 genes were selected for an orthogonal transcriptome screen (Fig. 2C, Table S4).

### A transcriptome-based secondary screening uncovers highly correlated phenotypes

We hypothesized that the phenotypes observed by microscopy might be reflected in their transcriptomes. Cells were transfected with siRNAs in triplicate and day 6 RNA samples were subjected to CEL-Seq2-based RNA-sequencing (Hashimshony et al., 2016). In addition to the 30 knockdowns, we also sequenced a day-by-day reprogramming time-course of control cells (Fig. 3A).

**Figure 3.**
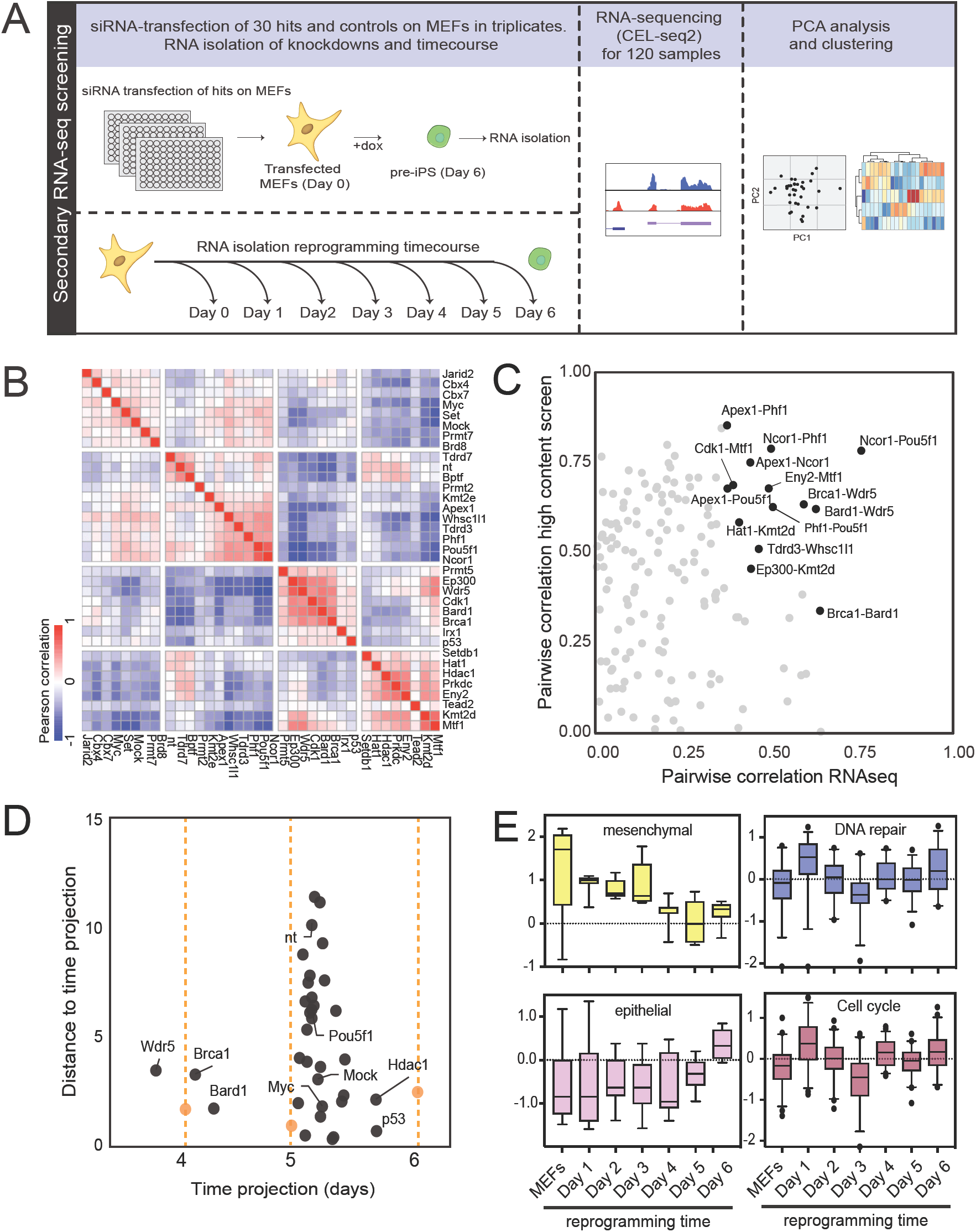
Transcriptome-based secondary screening. (**A**) Selected hits (30) and controls were transfected in triplicate and cultured until reprogramming day 6. The transcriptomes were analyzed together with a time series of control cells. (**B**) siRNA-to-siRNA Pearson correlation heatmaps based on transcriptomes. (**C**) Scatter plot representing pairwise siRNA correlations of transcriptomes (x-axis) and high-content image analysis (y-axis). siRNA pairs with highest correlations in both approaches are highlighted. (**D**) Analysis of the progression of reprogramming in knockdown cells compared to cells of the time course, based on PCA analysis of the transcriptomes and the projection of all data points on a curve fitted to the time course. (**E**) Boxplots representing log transformed and normalized gene expression values from the CELSeq2 time-course dataset. Each color depicts different groups of genes. (Experimental procedures, cf. Fig. S3 and Fig. S4).

We performed Principal Component Analysis (PCA) to the siRNA dataset for dimensionality reduction. The pairwise correlations between all the transcriptomes were calculated based on the top 200 transcripts associated with PC1 and PC2, and then clustered (Fig. 3B, left). We calculated similar pairwise correlations for the microscopy data, and identified gene pairs that correlated in both their colony phenotype and their transcriptome (Fig. 3C). The strongest correlations are observed between the Ncor1 - Oct4 pair and a triplet consisting of Wdr5, Brca1 and Bard1. Ncor1 was recently shown to physically interact with Myc and Oct4 (Zhuang et al., 2018), but the functional relationships between Wdr5, Brca1 and Bard1 were unknown. We performed siRNA deconvolution experiments measuring the number of Sall4-positive colonies of three independent siRNAs for Wdr5, Brca1 and Bard1 to exclude off-target effects. This analysis resulted in phenotypes similar to the pooled siRNAs in at least two out of three siRNA sequences with the same target (Fig. S3). In addition, high knockdown efficiencies of the Brca1, Bard1 and Wdr5 mRNA targets were verified at day 3 of reprogramming (Fig. S3).

As reprogramming is a dynamic process, we wondered how cells progress towards the iPSC state in each of the knockdown conditions. Notably, in PCA analysis, principal component 2 correlates strongly with time (r^2^ = 0.81; Fig. S4). To model the progression in each knockdown more precisely, we fitted a polynomial function to the time points and projected all other data on the time line by shortest distance (Fig. 3D, Experimental procedures). This distance reflects transcriptome changes that are unrelated to normal progression of reprogramming. Most siRNA knockdown transcriptomes, including the non-targeting (nt) and mock transfected controls have a transcriptome that is in between day 5 and day 6 of reprogramming, reflecting a mild non-specific effect of transfection. Silencing p53 and Hdac1 modestly speeds up reprogramming relative to nt controls (Fig. 3D). Three other genes, notably Wdr5, Brca1 and Bard1, show a strong delay in reprogramming with a short distance to the time projection of control cells. siWdr5 cells were comparable to normal cells between day 3 and 4, while siBard1 and siBrca1 were between day 4 and 5 (Fig. 3D). We analyzed our time series data to relate the early block observed with siWdr5, siBrca1 and siBard1 to known early reprogramming processes. The block is observed at the time of a major decrease of mesenchymal gene expression and preceding the activation of epithelial markers (Fig. 3E). For DNA repair and cell cycle genes there is an early wave of increased expression followed by downregulation, whereas random genes are stably expressed over the time course of reprogramming (Fig. S4). This time line raised the possibility that Wdr5, Brca1, Bard1 affect the repression of mesenchymal gene expression and the DNA damage response during early reprogramming. Moreover, based on the phenotypic and molecular co-correlation data we hypothesized that Wdr5, Brca1 and Bard1 functionally cooperate to control early stages of reprogramming.

### Brca1, Bard1 and Wdr5 functionally interact during early reprogramming

We asked whether *Wdr5, Bard1* and *Brca1* genes have similar expression dynamics during early reprogramming. Interestingly, the three genes follow a similar RT-qPCR profile, peaking in expression at day 3, and then slowly going down (Fig. 4A). To test the possibility of a functional interaction between these genes, the effect of their respective double knockdowns was measured and compared to the effect of the single knockdowns with regards to the number of pre-iPS colonies formed. All three single knockdowns displayed a significant reduction in number of Sall4-positive colonies, compared to the nt control (Fig. 4B). Therefore, the phenotypes of double and single knockdowns were calculated as the Sall4-positive colony ratio compared to the control. *Brca1-Bard1* double knockdown showed significantly more colonies than expected (Fig. 4C, left). This result was anticipated, as Brca1-Bard1 are well known physical interactors (Wu et al., 1996). Similarly, for both the *Wdr5-Brca1* and the *Wdr5-Bard1* double knockdowns, we also observed more colonies than expected, and this result was statistically significant for *Wdr5-Brca1* (Fig. 4C). To test whether Wdr5 is directly activating Brca1 and Bard1 gene expression, we determined the Brca1 and Bard1 expression levels after Wdr5 knockdown (Fig. 4D). Indeed, we find that this is the case at day 3, but also find that in response to either Bard1 or Brca1 depletion, Wdr5 expression was decreased. Taken together, *Brca1, Bard1* and *Wdr5* are co-expressed, mutually depend on each other, and interact functionally in reprogramming.

**Figure 4.**
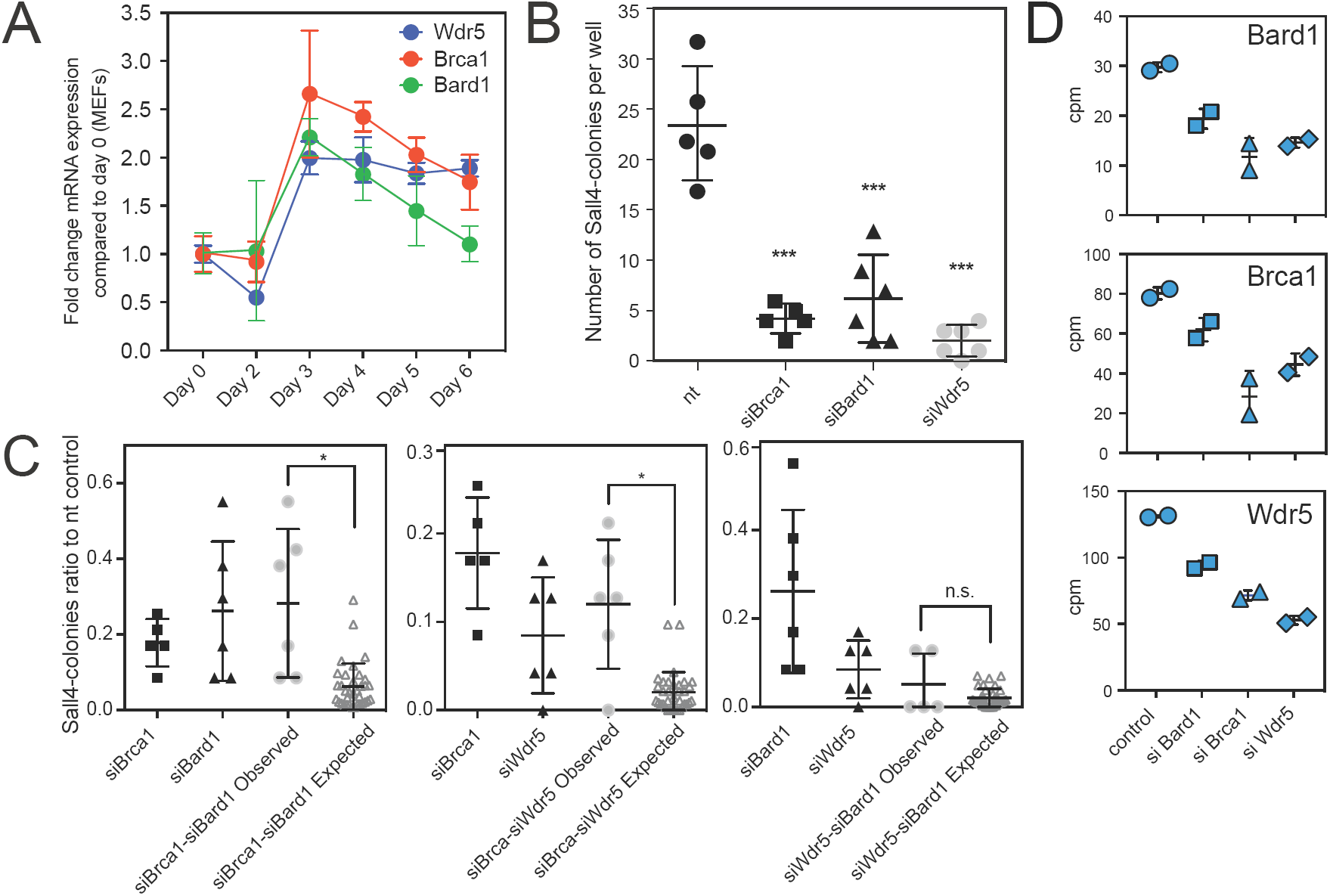
Brca1, Bard1 and Wdr5 functionally interact in early reprogramming. (**A**) Gene expression of Brca1, Bard1 and Wdr5 measured by RT-qPCR. Fold change was calculated relative to MEFs (day 0) gene expression. Each data point represents the mean value ± standard deviation of a biological duplicate. (**B**) Dot plot representing the number of Sall4-positive colonies measured by in-cell western in control and Brca1, Bard1 and Wdr5 knockdowns at day 6. Each dot represents one biological replicate and statistical significance determined by ANOVA is represented as *** p<0.0005. (**C**) Sall-4 colony ratios of the single and double knockdowns compared to the non-targeting (nt) control, measured by in-cell western. Functional interaction is determined by comparing the mean difference in double knockdown colony ratios: observed vs. expected. Each dot represents one biological replicate of an in-cell western for colonies at day 6 stained for Sall-4. Statistical significance p<0.05 (*) was calculated with two tailed T-test. **(D)** Dot plots to show Wdr5, Brca1 or Bard1 gene expression as counts per million reads (cpm) in siBard1, siBrca1, siWdr5 and nt control.

### Wdr5, Bard1 and Brca1 are functionally connected in the DNA damage response pathway

Brca1 and Bard1 have a known function in double strand break DNA repair. If Brca1 and Bard1 functionally interact with Wdr5, the prediction is that that all three knockdowns show an increase in DNA damage. The phosphorylated form of the histone variant H2A.X (γH2A.X) represents a reliable biomarker for DNA damage, because it is an immediate response upon the presence of double strand breaks (Sharma et al., 2012). Therefore, we employed FACS analysis to measure γH2A.X in the knockdowns (Fig. 5A-B). Reprogramming cells (nt control) showed a significant decrease in DNA damage response as compared to non-reprogramming MEFs, in agreement with literature showing that reprogramming resolves part of the DNA damage in somatic cells (Ocampo et al., 2016). Importantly, Wdr5 knockdown showed a significantly increased level of γH2A.X compared to the control (Fig. 5A). Nearly 90 % of the cells harbour γH2A.X in Wdr5 depleted cells (Fig. 5B, bottom panel). As expected, siBrca1 and siBard1 also showed a high percentage of γH2A.X positive cells (Fig. 5A, panels 2 and 3). To cross-validate our findings, we visualized γH2A.X by immunofluorescence. At day 3 of reprogramming, knockdown cells and controls were stained for either Oct4 (Fig. S5)or SSEA1 (Fig. 5C), and γH2A.X. In agreement with the results from the FACS analysis, nt control transfected reprogramming cells showed a decrease in γH2A.X compared to the MEFs. Depletion of Brca1, Bard1 or Wdr5 impairs the DNA damage response in reprogramming cells, resulting in more phosphorylated γH2A.X (Fig. 5, left and right). It should be noted that few cells or colonies show expression of SSEA1 in the siWdr5 cells, reflecting the early block of reprogramming progression (Fig. 3D).

**Figure 5.**
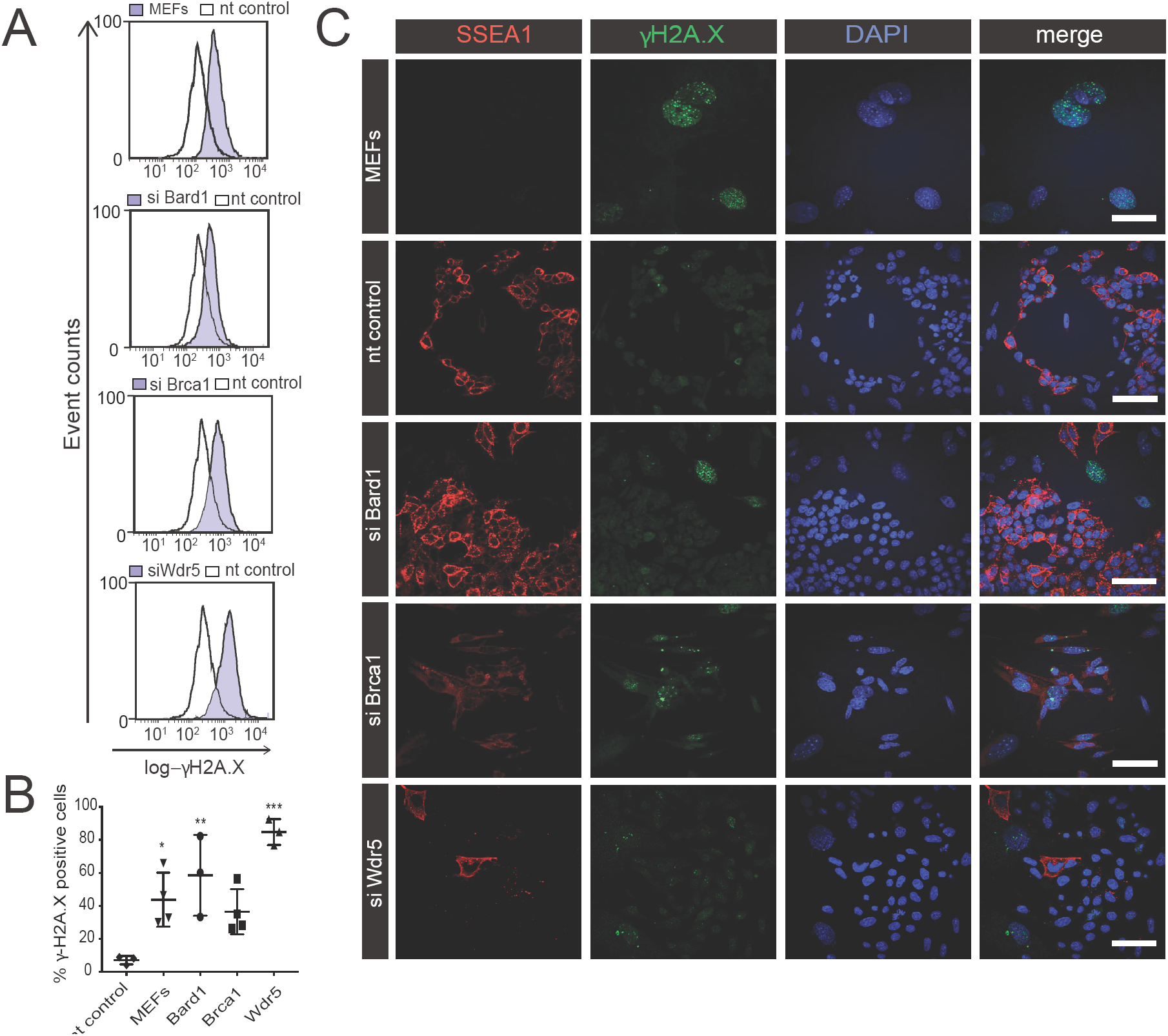
Wdr5, Bard1 and Brca1 are functionally connected in the DNA damage response pathway. (**A**) Representative FACS histograms showing the cell distribution with log-intensity of γH2AX in reprogramming populations measured in different conditions (white, nt; purple, siRNA). (**B**) Dotplot representing the quantification of γH2AX -positive cells in each condition in biological replicates. Each of the data points corresponds to a biological replicate, measured from independent experiments. Statistical significance was determined by one-way ANOVA. p<0.05(*), p<0.005(**) and p<0.0005(***). **(C)** Confocal images of reprogramming cells at day 3, stained for γH2AX (green) SSEA1 (red), counterstained with DAPI. Scale bar is 100 μm. See also Fig. S5.

### Wdr5, Brca1 and Bard1 are required for MET and DNA repair gene expression

Based on their timing of expression and the observed early block in reprogramming, we hypothesized that Wdr5, Brca1 and Bard1 also affect the MET. To test this hypothesis and to gain more insight into the Wdr5, Brca1 and Bard1 phenotypes, we performed deep RNA sequencing at day 3 and day 6 of reprogramming. We called differentially expressed genes and found 753, 1555, and 205 genes deregulated in respectively Wdr5, Brca1 and Bard1 knockdown cells following 3 days of OSKM induction (Fig. 6A). Wdr5, Brca1 and Bard1-depleted cells showed reduced expression of early pluripotency genes such as *Sall4, Cdh1* and *Epcam* (Fig. 6A).

**Figure 6.**
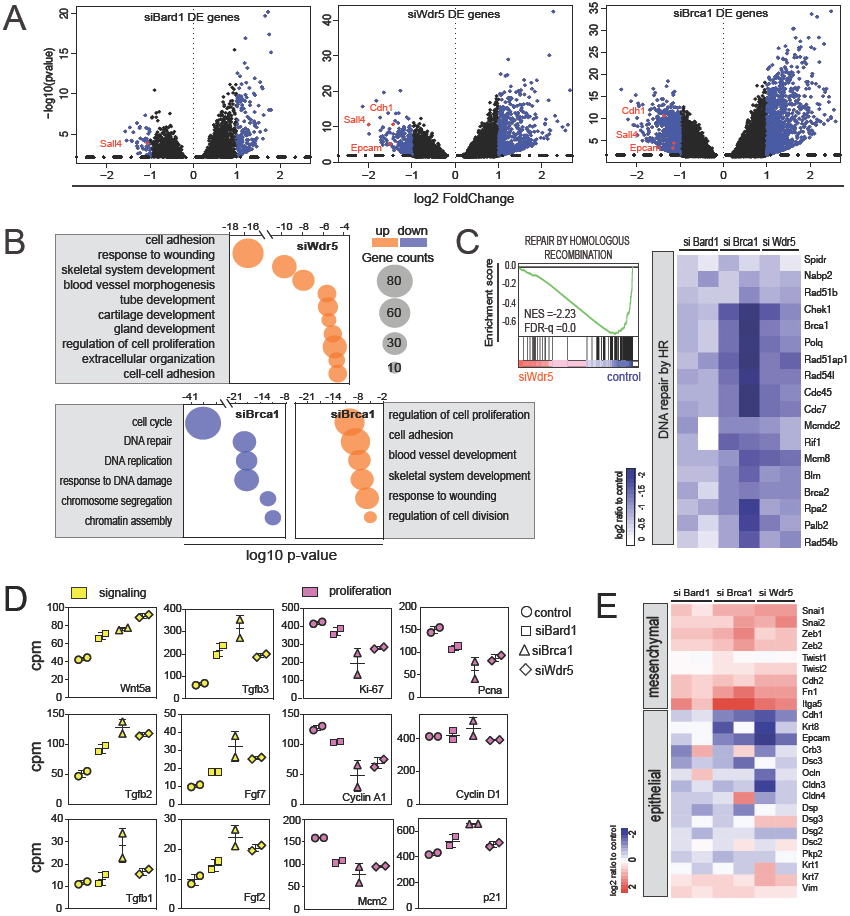
Wdr5, Brca1 and Bard1 depletion affects expression profiles of MET and DNA repair genes. (**A)** Volcano plots for siBard1 (left), siBrca1 (middle) and siWdr5 differential gene expression at reprogramming day 3. Blue highlight: differentially expressed genes (log2-fold change ≥1, adjusted p-value <0.05). (**B**) Bubble plot, showing examples of some of the most enriched terms (upregulated genes, orange; downregulated genes, blue) after Gene Ontology functional classification in siWdr5 and siBrca1 at Day 3. Bubble sizes represent the number of genes. (**C**) Gene Set Enrichment Analysis for DNA repair by homologous recombination (HR) comparing siWdr5 vs. control transcriptomes (left). Heatmap for siBrca1, siBard1 and siWdr5 samples showing DNA repair by HR genes, represented as log2-ratio relative to control (right) (**D**) Dot plots for signaling genes (magenta) and cell proliferation markers (yellow) quantified as counts per million reads (cpm) in control, siBrca1, siBard1 and siWdr5 cells. **(E)** Heatmap representing the log2-ratio of mesenchymal and epithelial gene expression of the three knockdowns relative to control. See also Fig. S6 and Table S5.

Differentially expressed genes in each knockdown were further probed for overrepresented gene ontology (GO) classes (Fig. 6B, Table S5). Brca1 knockdown causes a reduction in gene expression related to the cell cycle, response to DNA damage, and DNA repair (Fig. 6B). We asked whether the effects on the DNA damage response (Fig. 5) are reflected in the transcriptome of Wdr5 as well. To test this, DNA repair genes were probed in a Gene Set Enrichment Analysis (GSEA) (Mootha et al., 2003; Subramanian et al., 2005) comparing siWdr5 and control transcriptomes. Indeed, the negative normalized enrichment score (NES) indicated decreased expression of DNA repair genes in the siWdr5 as compared to the control (Fig. 6C, left). Furthermore, decreased expression of DNA repair genes in siWdr5 was similar to that of siBrca1 and siBard1(Fig. 6C, right and Fig.S6)

Wdr5 and Brca1 knockdowns shared a number of up regulated terms, including cell adhesion and developmental processes (e.g. skeleton or blood vessel development) (Fig. 6B). Regulation of cell proliferation is changed in Brca1, Bard1 and Wdr5 knockdowns; this GO term is enriched due to increased expression of *Tgfβ, Wnt, Bmp, Fgf* growth factors (Table S6, Fig. 6D). These growth factors decrease cell proliferation (Vega et al., 2004), but are also involved in epithelial to mesenchymal transitions (EMT) (Barrallo-Gimeno and Nieto, 2005), potentially counteracting the MET required for reprogramming. Several cell proliferation markers, such as *Pcna, Ki-67* and *Mcm2* were decreased in all three knockdowns, while *p21* (*Cdkn1a*) was up regulated (Fig. 6D). We assessed the gene expression levels of mesenchymal and epithelial markers in the three knockdowns and observed a clear increase in mesenchymal gene expression in the Wdr5, Brca1 and Bard1 knockdown cells relative to control cells (Fig. S6, Fig. 6E). Some epithelial genes were decreased (*Cdh1, Epcam* and *Krt8*), whereas others did not change substantially or were increased (Fig. 6E, Fig. S6).

Together, these data indicate that Wdr5, Brca1 and Bard1 not only cooperate in pluripotent colony formation (Fig. 4), but also share a functional interaction in the MET and the expression of DNA damage response genes during early reprogramming.

## DISCUSSION

This study reports on the colony morphology phenotypes of 300 chromatin-associated factors and on the transcriptome phenotypes of 30 factors during early reprogramming of fibroblasts towards induced pluripotency, constituting a highly relevant resource. Moreover, we have characterized the phenotypes involved in the mesenchymal-epithelial transition and the DNA damage response in more detail, and find cooperative contributions of three genes, Wdr5, Brca1 and Bard1 in both these processes.

Several complexes associated with the DNA damage response and replication are highly induced early in reprogramming (Hansson et al., 2012) and the p53-pathway is activated in cells harbouring substantial DNA damage (Marion et al., 2009). DNA damage may be associated with senescence, which can be rapidly induced by oxidative stress (Ben-Porath and Weinberg, 2005; d’Adda di Fagagna et al., 2003). It has been proposed that in vitro cell culture generates oxidative stress (Halliwell, 2003) and this could lead to an accelerated senescence (Parrinello et al., 2003). In agreement with this, low oxidizing conditions alleviate the reprogramming barrier imposed by senescence (Utikal et al., 2009). Some aging hallmarks, such as eroded telomeres (Lapasset et al., 2011; Marion and Blasco, 2010) and senescence-associated epigenetic marks (Ocampo et al., 2016) are reset by OSKM reprogramming. Brca1-Bard1 and Wdr5 may therefore alleviate a senescence-related block of reprogramming. In addition, the requirement of a DNA damage response could be related to the faster proliferation rates acquired early on in reprogramming (Polo et al., 2012; Ruiz et al., 2011). Embryonic stem cells, which proliferate in a similar fashion, require additional genome surveillance mechanisms to cope with fast DNA replication (Ahuja et al., 2016). The reduction in γH2A.X that we observe during normal reprogramming, however, is not common to all reprogramming systems (Gonzalez et al., 2013) and the observed differences could be due to presence of vitamin C in our medium, as the addition of antioxidants reduces genomic instability in reprogramming cultures (Ji et al., 2014).

We found a functional interaction of Wdr5, Brca1 and Bard1 in reprogramming. *Brca1, Bard1* could be direct or indirect targets of the SET/MLL complexes, of which Wdr5 is a subunit. In line with this possibility, ChIP analysis showed that Wdr5 binds regulatory regions of *Brca1, Bard1* and other genes involved in repair (Ang et al., 2011). Moreover, *Brca1* and *Bard1* transcripts are down regulated after silencing *Wdr5* (Fig. 4). In addition, not only are Brca1 and Bard1 involved in mitotic spindle organization and checkpoint gene regulation (Jin et al., 2009; Joukov et al., 2006; Wang et al., 2004), MLL/Wdr5 has been implicated in cell cycle regulation, mitotic progression and proper chromosome segregation (Ali et al., 2014; Liu et al., 2010; Ali et al., 2017).

Our study adds to the notion that colony morphology is linked to pluripotency (Abagnale et al., 2017; Kato et al., 2016; Narva et al., 2017) and is regulated by adhesion molecules, extracellular matrix and cytoskeleton forces. Upon differentiation, these processes orchestrate morphological changes such as loss of colony compaction, increase of cell area, colony flattening, together with changes in the pluripotency network (Narva et al., 2017). Therefore, colony morphology is a very important readout for reprogramming quality. Moreover, medium-high throughput screening of such multi-dimensional phenotypes is very powerful to identify functional interactions between genes. Brca1, Bard1 and Wdr5 depleted cells gave rise to fewer yet bigger, flat, symmetric colonies, due to a failure to properly down regulate mesenchymal cell adhesion molecules (Fig. 6). In addition, these cells fail to activate epithelial and early pluripotency genes. Our study links the DNA damage response to the MET program early in reprogramming through Brca1-Bard1 and Wdr5. Interestingly, the converse process of EMT may relate to DNA damage in kidney disease (Slaats et al., 2014) and cancer cells in culture (Chiba et al., 2012). Future work will further explore these relationships as well as gene-gene interactions that modify the phenotypical plasticity of reprogramming to induced pluripotency.

## EXPERIMENTAL PROCEDURES

### Data and Software Availability

Sequencing data are available at the GEO repository Superseries number GSE118680.

The code to reproduce reprogramming facilitator predictions by machine learning is available at https://github.com/simonvh/facilitators-penalosa-ruiz/. The code to reproduce the timeline projection is available at https://github.com/TimEVeenstra/Time-Curve-Projection/ (doi: 10.5281/zenodo.1405746).

### MEF Reprogramming and culture media

Passage 1-2 MEFs (mouse embryonic fibroblasts) were seeded at a density of 10,000 cells per cm^2^. Next day, MEFs were transduced at an MOI of 1 with Tet-STEMCCA lentivirus (Sommer et al., 2009), rtTA (Addgene#20342) and 8 μg·mL^-1^ polybrene. Next day (day 0), cells were transferred to either 1% gelatin-coated plates or mitotically inactive feeder cells, in reprogramming medium (Vidal et al., 2014).

### siRNA transfections and siRNA screenings

A custom Silencer siRNA library targeting around 300 mouse genes encoding chromatin factors was designed (Thermo Scientific/Ambion, Table S1) and distributed in 6 plates. Each gene in the library was targeted with three different siRNAs, which were pooled for transfection. For the high-content screening, the six pooled plates were transfected in quadruplicate. Every plate contained the following controls: siOct4 (siPou5f1), siMyc, siTrp53 and seven non-targeting (nt) controls. Reverse transfections in a 96-well plate format were performed as follows: 20 μL of transfection mix was prepared in each well before adding the cell suspension. This transfection mix consisted of 40 nM of pooled siRNAs, and 0.26 μL RNAiMAX lipofectamine (Thermo Scientific) diluted in Optimem (Thermo Scientific). After incubation for 10 minutes, 100 μL of cell suspension (3000-6000 cells) were added to each well. For transfections in a 6-well plate format, the protocol was scaled up accordingly. Before adding 1.8 mL cell suspension with 100,000 cells, 220 μL transfection mix was incubated in the wells for 10 minutes. The transfection mix consisted of 4 μL RNAiMAX and a final concentration of 40 nM siRNA, all diluted in Optimem.

### Immunostaining

Cells were cultured in 96-well Cell Carrier plates for microscopy (Perkin Elmer). After 6 days of reprogramming, cells were washed with PBS and fixed with 4% PFA for 15 min. After blocking and permeabilization, samples were incubated overnight with mouse anti-Cdh1 (Cell Signaling, 14472) and then with goat anti-mouse Alexa-488 for 2 hours. Staining with rabbit anti-Sall4 (Abcam, ab29112) was done overnight, followed by 3 hours incubation with goat anti-rabbit Alexa 568 and 40 μg·mL^-1^ DAPI. After antibody incubations, the cells were washed twice with PBS.

### High-content image acquisition and feature selection

Plates were imaged with an Opera High-content Screening System (Perkin Elmer) with a 4X air lens. Images were imported into the Columbus software platform (PerkinElmer). To segment colonies imaged on multiple z-planes, we used the maximum projection of z-planes. Sall4 staining was used to find and segment the colonies. Automated image analysis was used for image region segmentation and for extraction of shape and morphology features. Image regions touching the edge were removed. For more details, see Supplementary Information. After extracting all features for every plate from the automated pipeline, a Z-score normalization was applied per plate (Bakal et al., 2007) based on the mean values per feature. To select relevant features, a feature-to-feature Pearson correlation was calculated. Features with a high pairwise correlation (>0.8) were considered redundant.

### RNA sequencing and analysis

CEL-seq2 sample preparation (Hashimshony et al., 2016) was performed with a few adaptations (see Supplemental Information). Transcripts were mapped to *Mus musculus* genome version mm10 with Bowtie2 (Langmead and Salzberg, 2012), UMI corrected using standard settings of the CELseq2 pipeline (https://github.com/yanailab/CEL-Seq-pipeline), and matched to the gencode.vM13.annotation transcriptome. To relate knockdown data points to the progression of reprogramming, the transcriptomes were subjected to principal component analysis (PCA). Principal components 1 and 2 (PC1, PC2) were swapped (x-axis: PC2) and all data (knockdown and time series) were rotated 15 degrees. A second order polynomial curve was fitted to the time series (day 2-7), and all data points were projected on this curve (script: https://doi.org/10.5281/zenodo.1405747). For each data point, the projected x coordinate was used as a proxy for time, whereas the distance to the fitted time line (calculated using Pythagoras’ theorem) was used as a proxy for gene expression differences unrelated to the process of reprogramming. For normal RNA sequencing, Kapa-RNA HyperPrep kit with Ribo Erase was used for ribosomal depletion and library preparation (Roche, Kapa Biosystems), starting with 200 ng of total RNA. The libraries were amplified for 10 cycles, quantified with Qubit, checked for size distribution (300 bp) by Bioanalyzer (Agilent), and subjected to qPCR analysis before and after library preparation. Libraries were sequenced paired-end (Illumina NextSeq 500, read length 43 bp). Reads were aligned to the mouse genome (mm10) with STAR version 2.5.2b (Dobin et al., 2013).

### FACS analysis of DNA damage

Reprogramming MEFs were transfected with siRNAs in 6-well plates. After 3 days, cells were fixed on ice with 1 % PFA for 15 minutes and incubated with 70 % ice-cold ethanol at −20°C for two hours. Samples were then incubated with 100 μL mouse anti-phospho-H2AX (Millipore, diluted 1:100 in 0.25 % BSA 0.3 % triton/PBS) overnight at 4 °C. Then, cells were washed and stained with 100 μL Alexa 488 Goat anti-rabbit 488 (diluted 1:500) for 2 hours at room temperature. Finally, samples were incubated with propidium iodide (PI) overnight in the fridge and were sorted using an FC 500 (Beckman Coulter) machine. Data analysis was done with Flowing software v.2.5. As positive control, reprogramming MEFs were treated with 400 μg·mL^-1^ mitomycin C for 3 days.

### Double knockdowns and functional interactions

The observed Sall4-colony-ratio was calculated dividing the double-knockdown number of colonies by the average number of colonies of the control (6 biological replicates). The expected Sall4-colony ratio was calculated by multiplying the ratios of the single knockdowns (Mani et al., 2008). A P-value < 0.05 (two-tailed T-test) was considered significant. See supplementary information for details.

## AUTHOR CONTRIBUTIONS

Conceptualization GJCV, KWM, GPR; Methodology GPR, VB, GJCV, KWM, CB, JCRS; Experiments GPR, VB, JPG, SW, JVvV; Analysis GPR, VB, JPG, GJCV, SJvH, TEV; Writing the manuscript GPR, GJCV, KWM.

## ACKNOWLEDGEMENTS

The authors thank Dei M. Elurbe for useful suggestions, help on data analysis and processing of the sequencing files, Jessie A.G. van Buggenum for help with the colony counting script. Siebe van Genesen, Katie Tremble and E. Janssen-Megens for valuable technical help. VB and CB are funded by the Stand Up to Cancer campaign for Cancer Research UK, and Cancer Research UK Programme Foundation Award to C.B. (C37275/1A20146).

